# Combined agonism and antagonism of canonical and non-canonical progesterone receptors in triple-negative breast cancer cells potentiates cytotoxicity enhanced by PI3K inhibition

**DOI:** 10.64898/2026.06.25.734635

**Authors:** Paula E. Petrella, Jason W. Chen, Benjamin D. Cosgrove

## Abstract

Confounding the treatment options available to patients with triple-negative breast cancer (TNBC) are not only its purported lack of hormone receptor and growth factor receptor targets (ER^-^/ PR^-^/ HER2^-^), but its enrichment in plastic and chemoresistant breast cancer stem cells (BCSCs). Although descriptions of non-canonical PR expression in TNBC are rife in the literature, only canonical PR is considered in the definition of TNBC and is used to determine therapeutic strategy, not least because the utility of non-canonical PR modulation in TNBC chemoresistance is largely unexplored and poorly understood. Here we document the expression of three non-canonical PRs and the canonical PR (PGR) phosphorylated at Ser345 (p-PGR S345) in a panel of TNBC and luminal breast cancer cell lines, and employ combined PR agonists and antagonists to investigate the influence of PR activity on TNBC cell viability and PI3K inhibitor cytotoxicity. To examine the contributions of non-canonical membrane-associated PRs mPRβ and PGRMC1, we tested the agonist Org OD 02-0, a synthetic progestin targeted to mPRs; the PGRMC1 antagonist Ag-205; and the antagonist SPA70 against the cytosolic/nuclear PXR, in the background of pan-PI3K inhibition with Buparlisib (BUP). We also reveal that combinations of agonists and antagonists targeted to canonical and non-canonical PRs robustly potentiate the cytotoxic effects of PI3K inhibition, and also exhibit significant cytotoxicity on their own. Using functional assays, flow cytometry, immunocytochemistry and protein expression analyses, we found that simultaneously perturbing PRs and inhibiting PI3K function resulted in significantly greater cell death than vehicle control or BUP alone, and reduced the proportion of ALDH1^+^ BCSCs in two TNBC cell lines. We conclude that four types of PR are tractable targets in TNBC which participate in cell viability and enhance chemotherapy-induced cytotoxicity, and should be re-evaluated in an evolving definition of this challenging disease.

## Introduction

Effective treatment options for triple-negative breast cancer (TNBC), classically defined as estrogen and progesterone receptor negative and lacking in HER2 overexpression (ER^-^/PR^-^/HER2^-^), remain extremely limited, ostensibly due to not only its enrichment in chemoresistant breast cancer stem cells (BCSCs) ^1–3^ but also its insensitivity to standard-of-care hormone and HER2 receptor-targeted chemotherapies, including Tamoxifen and Herceptin respectively. Prior reports, however, detail expression of functional variants of the canonical nuclear progesterone receptor PR (PGR) in TNBC ^4^; as well as membrane progesterone receptors (mPRs) ^5,6^, progesterone receptor membrane component 1 (PGRMC1) ^7,8^; and the pregnane X receptor PXR (PXR) ^9,10^, and here we report the expression of phospho-PGR S345 (p-PGR S345) as well. While the existence of these receptors in TNBC cells do not necessarily imply vulnerability to hormone-based therapy, their potential in this capacity is compelling and little explored.

To investigate the effects of PR modulation on TNBC chemoresistance, we tested agonism and antagonism of canonical and non-canonical PRs in combination with Buparlisib (BUP), a pan-PI3K inhibitor, in three TNBC cell lines (BT-20, HCC1937 and MDA-MB-231) and a luminal-A BC line with constitutive expression of PGR (T47D) (Table 1). Both progesterone (P4) and PI3K are involved in the maintenance and viability of cancer stem cells (CSCs), with P4 participating in the generation and regulation of CSCs in both BC and basal-like non-transformed mammary cells ^11–16^ while oncogenic PI3K activation is associated with BCSC viability and maintenance ^17–21^. Notably, mifepristone (MIF), a PR antagonist, suppresses BCSCs in TNBC ^22^. Furthermore, P4 activates PI3K through its cognate receptors ^7,12,23^. In addition to its role as a transcription factor, canonical PGR associates with PI3K and participates in the activation of PI3K and MAPK signaling pathways ^24–33^. Interestingly, P4 was recently shown to rescue zebrafish development abrogated with leflunomide-induced nucleotide stress ^34^, a stress similar to that caused by PI3K inhibition ^35^ which we explored previously in TNBC ^36^. As leflunomide is also a PXR antagonist, this finding suggests a therapeutic opportunity in combining PR modulation with PI3K inhibition. Historically, the lack of TNBC therapeutic response to PGR-targeted treatments may not have resulted solely from the scarcity of canonical PGR expression but may turn on the additional targeted perturbations of mPRs, PGRMC1 and PXR.

**Table 1.** Model human triple-negative and luminal breast cancer cell lines used in this study. Tumor subtypes and commonly mutated genes and loci in each cell line used in this study. Reference data from Sanger Institute (cellmodelpassports.sanger.ac.uk), Cellosaurus at SIB Swiss Institute of Bioinformatics (cellosaurus.com), and ATCC (ATCC.org).

| CELL LINE | TUMOR SUBTYPE | HISTOLOGY | TUMOR SOURCE | COMMONLY MUTATED GENES | MUTATED LOCI (cDNA / protein) |
| --- | --- | --- | --- | --- | --- |
| BT-20 | Unclassified | Carcinoma | Primary | *CDKN2A* - homozygous<br>*PIK3CA* - heterozygous<br>*PIK3CA* – heterozygous<br>*RB1* - heterozygous<br>*TP53* - homozygous | c.1_471del471 / p.0?<br>c.1616C>G / p.Pro539Arg<br>c.3140A>G / p.His1047Arg<br>c.1544C>T / p.Pro515Leu<br>c.394A>C / p.Lys132Gln |
| HCC1937 | Basal-like 1 | Ductal carcinoma | Primary | *BRCA1* - homozygous<br>*PTEN* - homozygous<br>*RB1* - homozygous<br>*TP53* - homozygous | c.5266_5267insC / p.Gln1756Profs*74<br>Deletion<br>c.2212_2325del114 / p.Thr738_Arg775del38<br>c.916C>T / p.Arg306Ter |
| MDA-MB-231 | Mesenchymal stem-like | Adenocarcinoma | Metastasis, pleural effusion | *BRAF* - heterozygous<br>*CDKN2A* – homozygous<br>*KRAS* - heterozygous<br>*NF2* - homozygous<br>*TP53* - homozygous | c.1391G>T / p.Gly464Val<br>c.1_471del471 / p.0?<br>c.38G>A / p.Gly13Asp<br>c.691G>T / p.Glu231*<br>c.839G>A / p.Arg280Lys |
| T47D | Luminal A | Ductal carcinoma | Metastasis, pleural effusion | *PIK3CA* - heterozygous<br>*TP53* - homozygous | c.3140A>G / p.His1047Arg<br>c.580C>T / p.Leu194Phe |

P4 is an agonist for all four classes of PR, and MIF also interacts with all four in a broad range of contexts ^37–43^. For selective mPRβ agonism, we employed a synthetic progestin, Org OD-02-0 ^5,44^ and use the PGRMC1 and PXR antagonists Ag-205 ^7,45,46^ and SPA70 ^47,48^ respectively. Through targeted simultaneous agonism and antagonism of these PRs in combination with BUP, we discovered treatment combinations which significantly increase cytotoxicity caused by PI3K inhibition (PI3Ki), and also found therapeutic synergy between agonist and antagonist pairings in the absence of PI3Ki. Specific treatment combinations also elevated stress and survival signaling in multiple cell lines. Intriguingly, we found that combined P4 and PI3Ki treatment significantly reduced the fraction of ALDH1^+^ BSCSs below that of BUP treatment alone in BT-20 cells and below vehicle control (VC) in HCC1937 cells, indicating a defense against the adaptive phenomenon of enriched ALDH1^+^ BCSCs following chemotherapeutic treatment^49,50^.

P4-responsive receptors with structures and functions as disparate and pleiotropic as these four pose a broad spectrum of opportunities for targeted chemotherapeutics in TNBC and other cancers. The future of TNBC treatment may lie in a more nuanced application of PR-targeted therapies, with a new definition of TNBC emerging that encompasses shades of gray, as do recent investigations into HER2-associated BCs ^51,52^ and ER-negativity in TNBC ^53^ and BC ^54^. Our approach highlights the advantages in modulating PR activity in the treatment of TNBC with small molecule chemotherapy and underscores the utility of reassessing the dogma of triple-negativity.

## Results

### Four types of progesterone receptors are expressed in multiple cellular compartments in TNBC and luminal BC cell lines

First, we sought to examine the expression and localization of progesterone receptor proteins in TNBC cell lines. We selected three TNBC cell lines, BT-20, HCC1937, and MDA-MB-231, and the T47D cell line which is a PR^+^ luminal A BC subtype, all of which are well-characterized and exhibit a variety of mutational profiles (see Table 1 for more detail). We cultured these four BC cell lines in complete medium for 72h and performed immunocytochemistry, staining them for PGR, p-PGR S345, mPRβ, PGRMC1, or PXR (Figure 1). All cell lines exhibited >97% positivity for nuclear localization of p-PGR S345, suggesting frequent ligand-activated PGR (Figure 1A, E, I, M; quantifications by cell line in 1R-U). As expected, T47D cells exhibited stronger nuclear expression of PGR (>30%) than TNBC cells (<10%; data not shown) (Figure 1M, U). mPRβ was found to be expressed in >97% of cells in all four cell lines, in the plasma membrane and in the perinuclear compartment, likely associated with the Golgi apparatus (Figure 1B, F, J, N) ^55,56^. PGRMC1 was found to occupy not only the cell membrane, but also the nuclear and perinuclear regions in >95% of cells across all cell lines (Figure 1C, G, K, O), as it is known to be expressed in the endoplasmic reticulum and nucleoli of human cells^56^. PXR has been shown to reside in the cytoplasm, translocating to the nucleus upon ligand binding ^57^; we found it to occupy the nuclear and cytoplasmic compartments in >95% of cells in all cell lines (Figure 1 D, H, L, P). These observations document that activated PGR, mPRβ, PGRMC1, and PXR all are expressed in various TNBC and luminal A cell lines. For quantifications of ICC images (Figure 1R-U), PGR and p-PGR S345 were counted per nucleus and all other PRs were tabulated per cell body. Of note, all cell lines appeared to express all PRs, except canonical PGR, at near 100% positivity, with count reductions primarily for mitotic, apoptotic and multi-nucleated cells.

**Figure 1.**
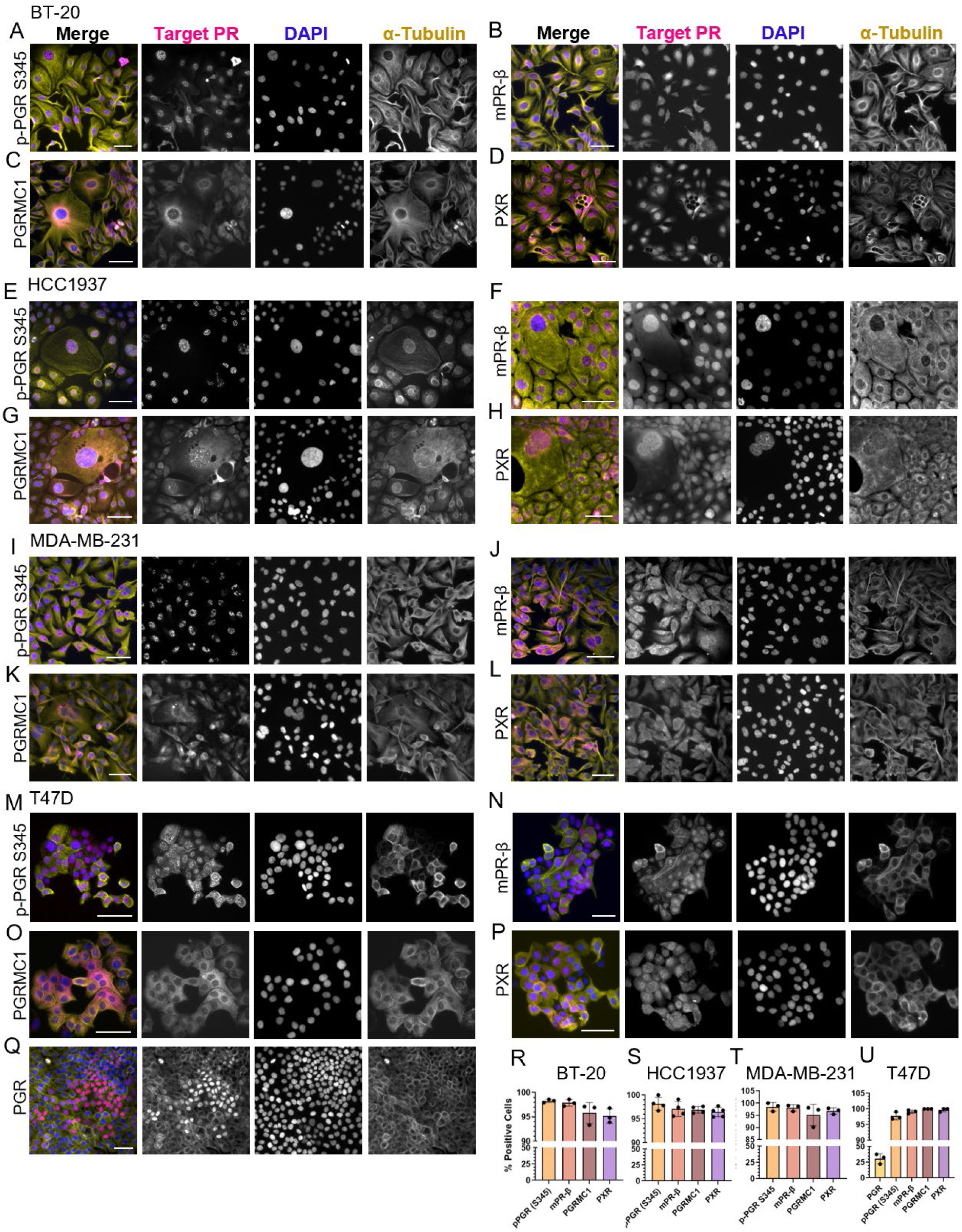
Four types of progesterone receptors are expressed in various cellular compartments in TNBC and luminal BC cell lines. TNBC and luminal BC cell lines were treated for 72h with Vehicle Control (VC), then fixed and stained with the appropriate anti-progesterone receptor (PR) antibodies and appropriate fluorophore-conjugated secondary antibodies, with α-tubulin and DAPI counterstaining. BT-20 cells immunostained for (A) Phospho-PR (pS345); (B) mPRβ; (C) PGRMC1; (D) PXR; and, respectively, HCC1937 in (E), (F), (G), (H); MDA-MB-231 in (I), (J), (K) and (L); and T47D in (M), (N), (O), (P) with PGR in (Q). Quantifications of PR-positive cells in each cell line in (R-U). Bar graphs report mean ± SD of n=3-4 replicates. Scale bars, 100 μm.

To examine the effects on PR expression of phenol red, a suspected estrogen mimic ^58,59^, and standard FBS containing lipids and steroids, we grew separate cohorts of BT-20 cells in growth media and FBS as well as in phenol red-free media containing charcoal-stripped FBS. We compared ICC and viability results using cells from each media formulation and found insignificant differences between the two (data not shown).

### Canonical and non-canonical progesterone receptor proteins are expressed at baseline in TNBC and luminal BC cell lines

To examine whether canonical and non-canonical PR protein expression is detectible at baseline in TNBC and luminal A cell lines, we performed immunoblotting after culturing cells for 72h in complete medium (Figure 2). We detected expression of p-PGR S345, mPRβ and PGRMC1 in all cell lines, and PXR in TNBC cells only (data not shown) (Figure 2A, B). For quantification, expression data was normalized to housekeeping genes GAPDH, α-tubulin or α-actinin as appropriate (1C-F). Our results indicate TNBC and luminal A BC cell lines express canonical and non-canonical progesterone receptors in normal growth conditions.

**Figure 2.**
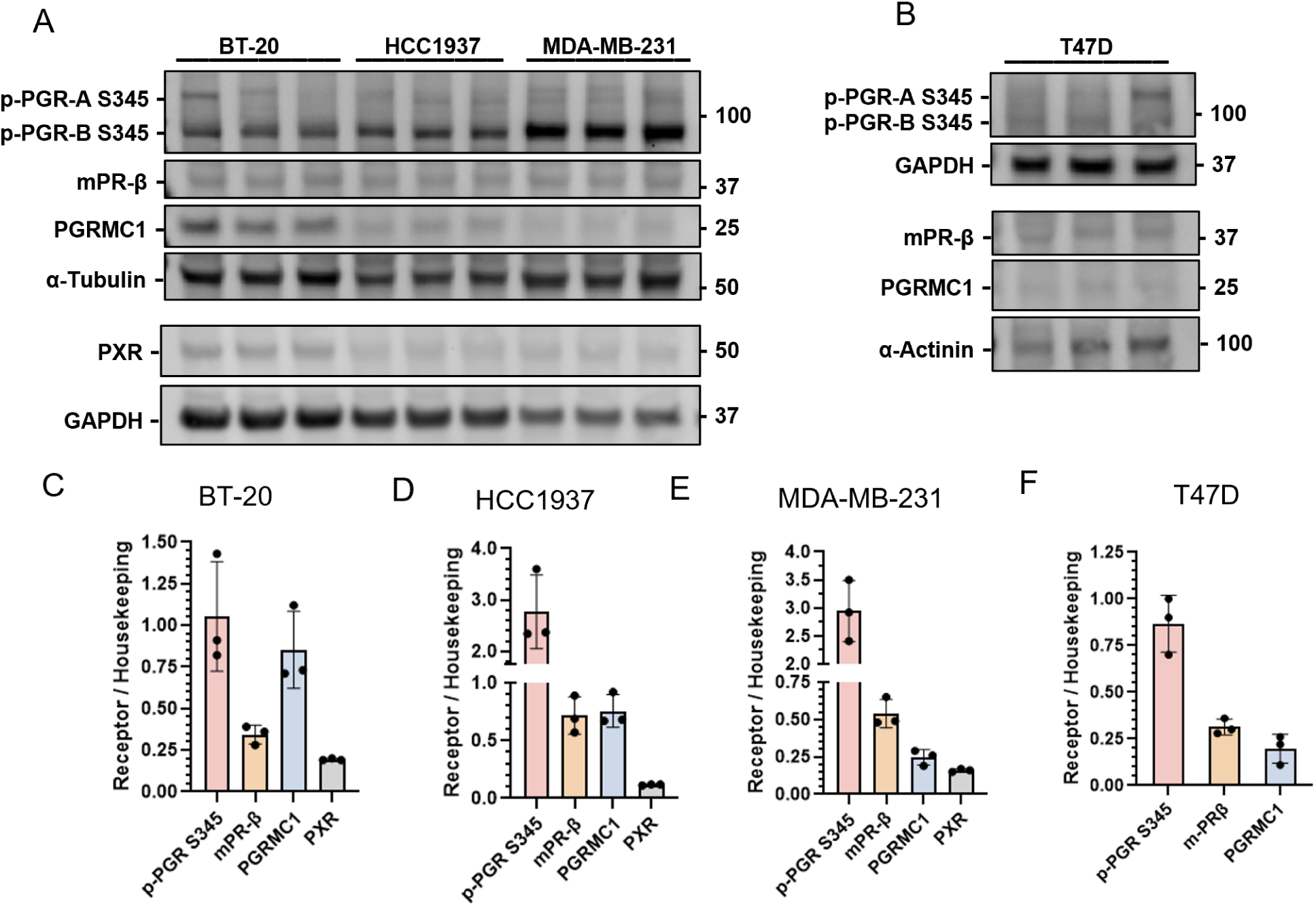
Canonical and non-canonical progesterone receptors are expressed at baseline in TNBC and luminal breast cancer cell lines. Western blots detailing expression of progesterone receptors phospho-PGR A/B S345 (p-PGR S345), mPRβ, PGRMC1 and PXR in TNBC cells BT-20, HCC1937 and MDA-MB-231 (A) or luminal A BC cells T47D (B). Cells were grown in complete medium for 72h. Lysates probed with the indicated antibodies. PXR not detected in T47D (data not shown). Quantifications for BT-20 in (C); HCC1937 in (D); MDA-MB-231 in (E); T47D in (F); bar graphs report mean ± SD of n=3 replicates. Analytes p-PGR S345, mPRβ and PGRMC1 in BT-20, HCC1937 and MDA-MB-231 cells normalized to α-Tubulin in (A); PXR in these cells normalized to GAPDH (A). In T47D cells, p-PGR S345 was normalized to GAPDH; mPRβ and PGRMC1 were normalized to α-Actinin (B).

### Co-treatment with progesterone and mifepristone induces cell death and potentiates the chemotherapeutic effect of the pan-PI3K inhibitor Buparlisib in TNBC and luminal BC cells

Given our observed PR expression in TNBC and luminal BC cell lines in Figures 1 and 2, and the ability of P4 to rescue zebrafish development stalled by leflunomide, a pyrimidine synthesis inhibitor and PXR antagonist ^34^, we hypothesized that activating and/or inhibiting PR function may influence PI3K inhibition outcomes in TNBC cells. First, we focused on a chemotherapeutic challenge model with the pan-PI3K inhibitor Buparlisib, which exhibits partial chemotherapeutic success at moderate doses (∼0.5-2.0 μM) which we previously attributed to an increase in residual ALDH1^+^ cancer stem-like cells ^36^. Here we tested co-treatment of BT-20, HCC1937, MDA-MB-231 TNBC cells and T47D luminal BC cells for 72h with BUP and an array of dosing combinations of progesterone (P4; an agonist for canonical PGR and non-canonical mPRβ, PGRMC1, and PXR) and/or mifepristone (MIF; a broad antagonist of these four PRs) (Figure 3). We analyzed 1.0 μM BUP in BT-20 cells because of their exquisite sensitivity, versus 2.0 μM BUP in all other cells lines. As expected, we observed that the two *PIK3CA* mutant cell lines, BT-20 and T47D, exhibited strongly reduced cell viability responses to BUP alone (Figure 3A-D, M-P), with the other two TNBC lines showing lower sensitivity (Figure 3E-L). We noted a broad pattern across all cell line responses to treatment, in that both BUP alone, and the combination of P4+MIF elicited significant cell death versus VC (Figure 3D, H, L, P, Q). In every cell line, P4+BUP and P4+MIF+BUP induced additional significant cell death beyond BUP alone, as did MIF+BUP in all but BT-20 cells (Figure 3D, Q). Hence, across cell lines, combined treatments with P4+MIF in itself significantly reduced cell viability, and also significantly potentiated the effects of BUP. In BT-20 cells for example (Figure 3C-D), P4+MIF reduced cell viability to 68.2%; BUP treatment alone resulted in 33.7% cell survival; and the addition of P4+MIF to BUP reduced that number to 14.8% (all p < 0.01). The HCC1937 cells exhibited a more drastic reduction in cell numbers with BUP alone (65% viable) than with P4+MIF alone (87.6%) (both p < 0.01; Figure 3G, H, Q), but P4+MIF+BUP elicited significantly lower viability than BUP, at 56% (p < 0.05). MDA-MB-231 cells exhibited roughly equal cell death in response to P4+MIF or BUP alone (80.1% and 81.5% viability respectively, both p < 0.01), (Figure 3K, L, Q). Similarly to the HCC1937 cells, T47D cells were significantly reduced by both BUP alone, at 44% viability and P4+MIF at 57.8% viability, and these results were improved upon by combining all three drugs, which resulted in a significant reduction beyond BUP alone, to 28.4% viability (all p < 0.01; Fig 3O, P, Q). These observations suggest that simultaneous agonism and antagonism of broad types of PRs in BCs can restrict proliferation as well as potentiate PI3K chemotherapeutics in a cell line-specific manner.

**Figure 3.**
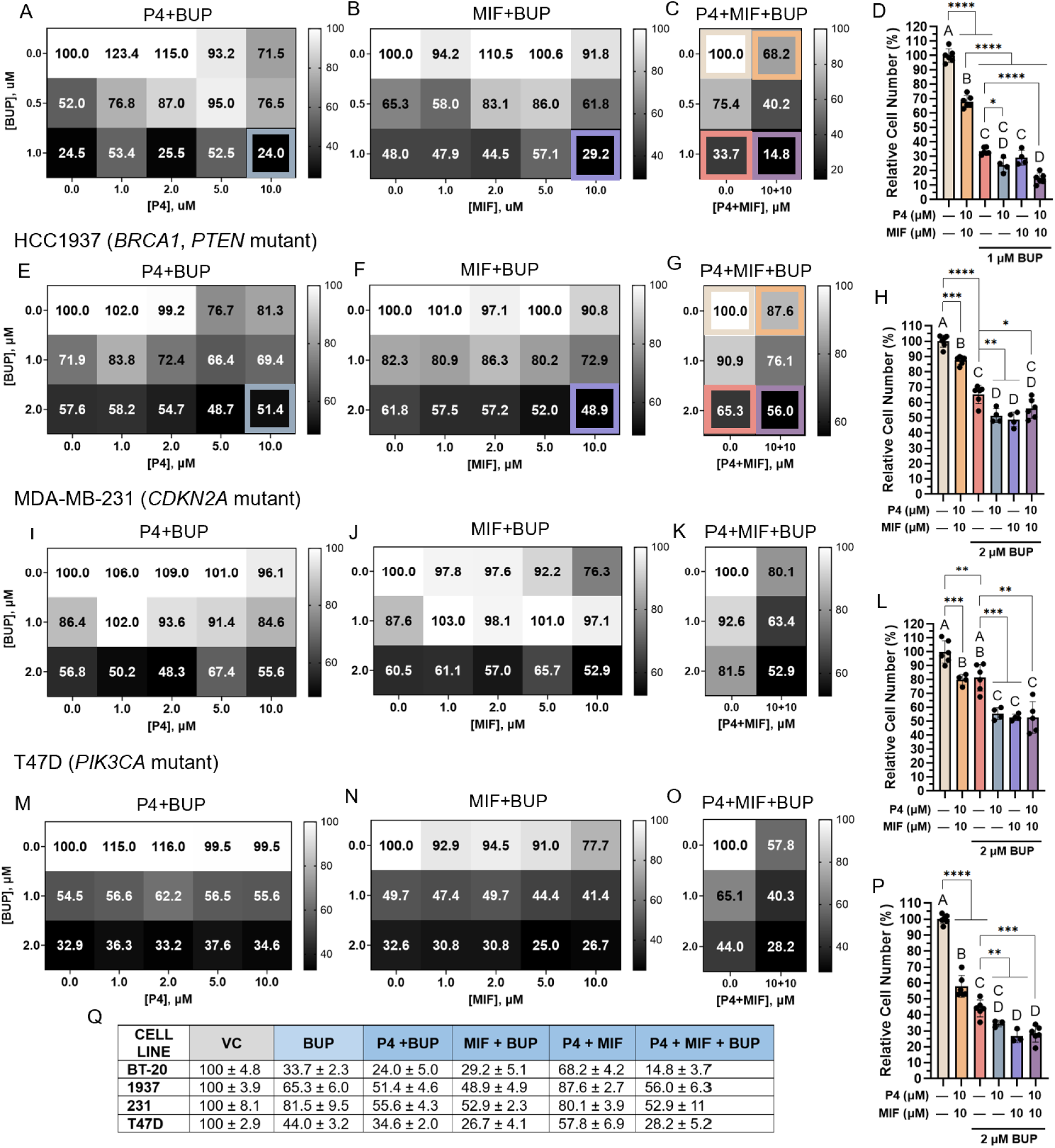
Co-treatment with progesterone and mifepristone induces cell death and potentiates the chemotherapeutic effect of the pan-PI3K inhibitor Buparlisib in TNBC and luminal BC cells. TNBC cells BT-20 (A-D), HCC1937 (E-H) and MDA-MB-231 (I-L) and PR^+^ luminal A BC subtype T47D cells (M-P) were treated for 72h with the phosphoinositide 3-kinase (PI3K) inhibitor Buparlisib (BUP; 0-1 μM in BT-20 cells; 0-2 μM in all others), and/or progesterone (P4; 0-10 μM) and mifepristone (MIF; 0-10 μM) in various dosing combinations, and relative cell viability was measured. Heatmaps report the average of n=3-6 replicates, and use color-scales unique to each. Bar graphs report mean ±SD of n=3-6 replicates. Data are reported as cell number relative to vehicle control (VC). Summary table in Q indicates % viability ± SD; VC data from C, G, K, O for BT-20, HCC1937, MDA-MB-231 and T47D respectively. Conditions that do not share a letter designation are significantly different. Welch’s t-test, * p < 0.05, ** p < 0.005, *** p < 0.0005, **** p < 0.0001.

### Simultaneous perturbation of estrogen and progesterone receptors restricts TNBC cell viability when combined with PI3K inhibition

Estrogen receptors (ERs) are also expressed in TNBC ^60–64^ and are known to interact with canonical and non-canonical PRs ^65–68^. Because the human *PGR* gene contains estrogen response elements and E2 can induce the expression of *PGR*, ^66,67,69^ we inferred that perturbing estrogen receptor signaling might also affect PR-based chemotherapeutic benefits. To test whether indirect agonism of PGR with estradiol (E2), an ER agonist, combined with PGR antagonism, causes TNBC cell death, we examined BT-20 cell viability under 72h treatment with 0.34 μM of E2, 0.0-1.0 μM BUP, and 10 μM of either or both P4 and MIF (Figure S1A, C). We observed that E2+MIF caused a significant reduction in cell viability versus VC (66.0% viable, p < 0.01), followed by E2+P4+MIF at 82.6% viable (p < 0.05). When combined with 0.5 μM BUP, E2+MIF significantly reduced viable cells to 63.6% versus BUP alone at 82.8% (p < 0.01). Similarly, E2+P4+MIF in combination with 0.5 μM BUP caused a significant decrease in cell viability, to 66.9% versus BUP (p < 0.01). Upon combination with 1.0 μM BUP, significant cell death was observed versus BUP alone (70.3%) with E2+P4 (59.2% viable, p < 0.05), E2+MIF (47.1%) and E2+P4+MIF (54.0%) (both p < 0.01). These effects echo the cytotoxic benefits we found when combining PR agonism and antagonism in TNBC.

To investigate the effects of ER antagonism combined with PR agonism and/or antagonism in the background of PI3K inhibition, we tested BT-20 cell viability under 72h treatment with 2.0 μM of fulvestrant (FULV), a selective estrogen receptor degrader with no known agonist effects ^70^, 0.0-1.0 μM BUP, and 10 μM of either or both P4 and MIF (Figure S1B, D). On its own, FULV had minimal effects on cell viability (98.1%, ns), but when combined with P4 (84.0% viability, p < 0.05), MIF (58.8% viability) or both (51.8% viability), cell viability was significantly reduced from VC (both p < 0.01). When added to 0.5 μM BUP, all treatments containing FULV induced greater cell death than BUP alone at 96.5% viability (all p < 0.01). Similarly, all treatments combined with 1.0 μM BUP (70.2%) resulted in significant cell death except FULV alone (71.1%, ns; all others p < 0.01). These unexpected cytotoxic effects of ER agonism and antagonism in combination with PR agonism and antagonism mirror our findings with PR perturbations, and provide evidence of hormone receptor treatments potentiating the cytotoxic effects of PI3K inhibition in TNBC.

### Progesterone receptor perturbations in combination with PI3K inhibition induce stress and survival adaptations in TNBC cells

To investigate the effects of BUP (B), P4 (P) and MIF (M) treatments on TNBC cell stress and survival signaling as compared to VC, we performed Western blotting on HCC1937 cell lysates after 72h drug treatment, to examine changes in p-p38, MDR1, PARP and cleaved PARP expression, as well as effects on the PI3K and MAPK signaling pathways via p-Akt and p-ERK expression respectively (Figure 4). All treatments with BUP (Figure 4A, D, G, H) drastically reduced p-Akt expression as expected (p < 0.05, Figure 4B). Akt expression, which can increase during stress as a compensatory survival response ^71^, rose above VC in all treatment conditions, reaching its highest levels in P+B and P+M+B conditions (Figure 4E). The expression of p-ERK 1/2, conversely, reached its lowest points with P+B and M+B treatments (Figure 4C), which may indicate treatment efficacy ^72^. PARP expression, which can increase during treatment evasion and resistance to apoptosis ^73^, rose significantly from VC to M+B and P+M+B treatments (both p < 0.05; Figure 4I). Cleaved PARP expression, an early indicator of apoptosis, followed a different pattern, declining from VC through BUP and P+B treatment, and rising again in M+B and P+M+B conditions (Figure 4J). Stress signaling via the p-JNK 1/2 pathway rose significantly higher in the P+M+B treatment condition than in BUP treatment (p < 0.01; Figure 4K). Conversely, p-p38 signaling increased with BUP treatment compared to VC, but fell below VC in all other conditions (Figure 4F). Expression of the drug efflux pump MDR1 was starkly highest in the P+B condition, significantly above VC, BUP and MIF+BUP treatments (all p < 0.05; Figure 4M). Taken together, these data indicate combined PR and PI3K perturbations increase stress signaling and treatment adaptation in TNBC cells.

**Figure 4.**
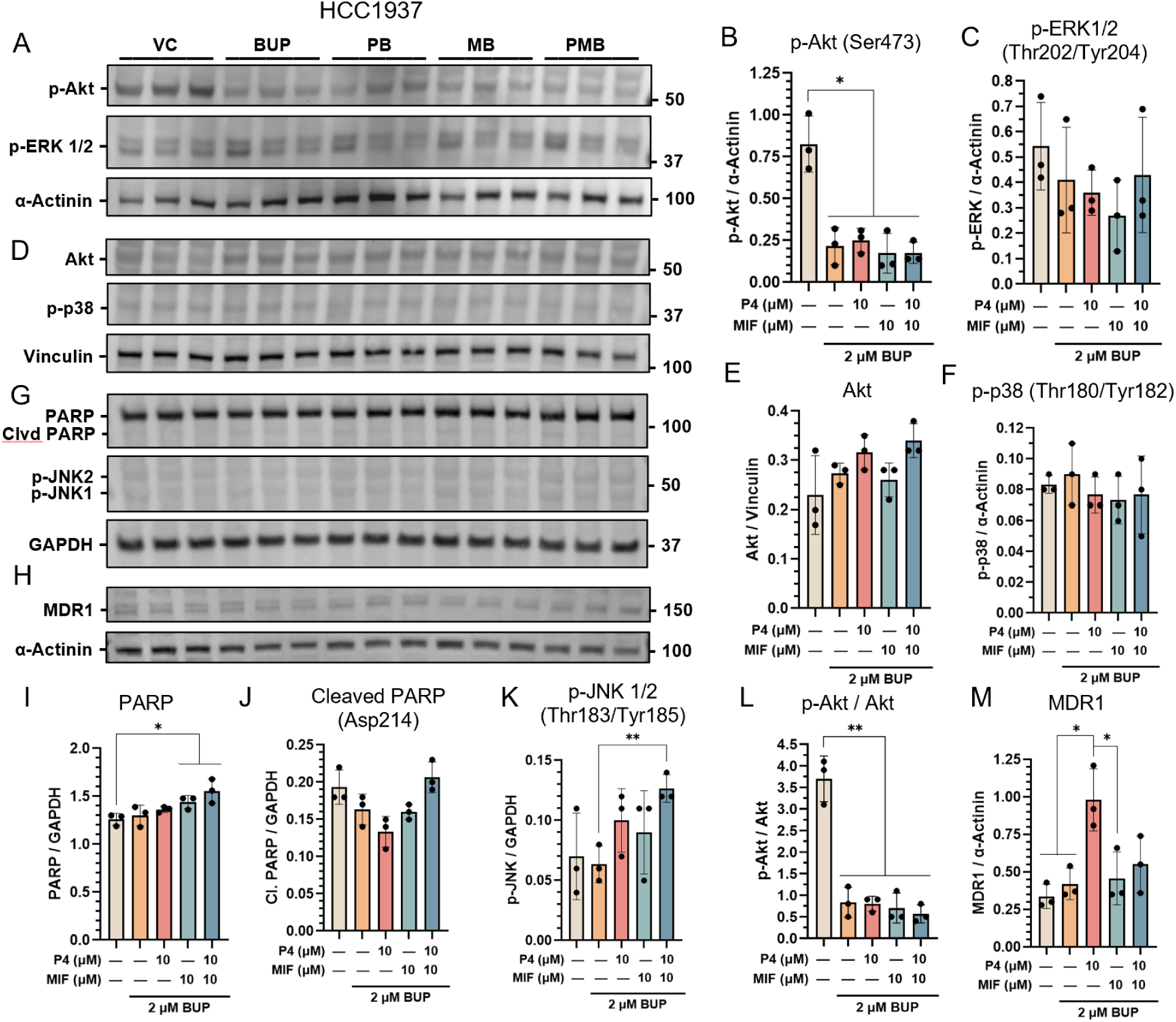
Progesterone receptor perturbations in combination with PI3K inhibition induce stress and survival adaptations in TNBC cells. Western blots (A, D, G, H) and quantifications (B, C, E, F, I-M) detailing protein expression following 72h treatment of HCC1937 cells with vehicle control (VC), Buparlisib at 2 μΜ (B; BUP) alone or in combination with progesterone (P) and/or mifepristone (M) both 10 μΜ. In (A) phospho-Akt (Ser473) and phospho-ERK 1/2 (Thr202/Tyr204) were probed; quantifications in (B, C); this blot was re-probed in (D) for Akt and phospho-p38 (Thr180/Tyr182); quantifications in (E, F) and p-Akt/Akt quantified in (L). PARP, cleaved PARP (Asp214) and phospho-JNK 1/2 (Thr183/Tyr185) were probed in (G); quantifications in (I-K); MDR1 was probed in (H); quantification (M). Bar graphs report mean ± SD of n=3 replicates; data normalized to housekeeping genes α-actinin (A, H), vinculin (D), or GAPDH (G). Welch’s t-test, * p < 0.05, ** p < 0.005.

### Individual and combined agonism and antagonism targeted to non-canonical PRs in TNBC and luminal BC cells induce cell death and potentiate PI3K inhibition

To gain a deeper understanding of the PR mechanisms driving these combinatorial treatment benefits, we first examined the relative transcript expression levels (TPM normalized) of *PGR* (encoding PGR A/B), *PAQR8* (encoding mPRβ), *PGRMC1*, and *NR1I2* (encoding PXR) for all four cell lines in the DepMap Consortium Expression Public 23Q4 database ^74^ (Figure 5K). As expected, we found that *PGR* transcripts are detected in greater abundance in T47D cells than in any TNBC line. Notably, *PAQR8* transcripts are most abundant in MDA-MB-231 and T47D cells; *PGRMC1* transcripts are detected in all four lines; and *NR1I2* is most highly expressed in HCC1937 cells. Hence, we focused on the cell lines that most highly expressed each PR.

**Figure 5.**
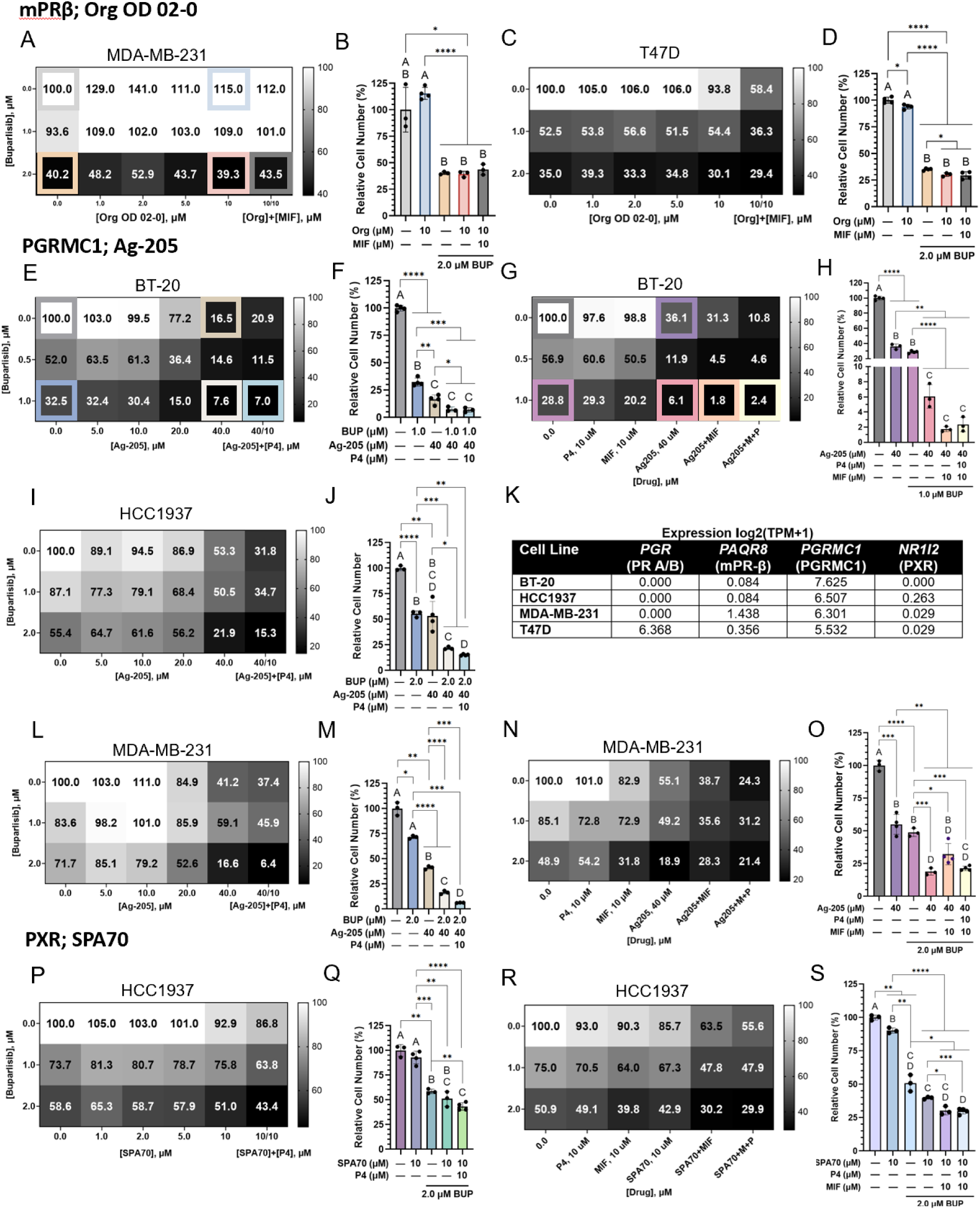
Individual and combined agonism and antagonism targeted to non-canonical PRs in TNBC and luminal BC cells induce cell death and potentiate PI3K inhibition. Viability heatmaps and quantifications for TNBC and luminal BC cell lines treated for 72h with: mPRβ agonist Org OD 02-0 (Org) at 0-10 μM, PI3K inhibitor Buparlisib (BUP) at 0-2 μM, and/or mifepristone (MIF; 0-10 μM) (top panels, A-D); PGRMC1 antagonist Ag-205 at 0-40 μM; BUP 0-1 μM in BT-20 cells, 0-2 μM in all others, and/or P4 0-10 μM and/or MIF 0-10 μM (middle three panels, E-J, L-O); and PXR antagonist SPA70 at 0-10 μM, BUP 0-2 μM, and/or P4 0-10 μM and/or MIF 0-10 μM (bottom panels, P-S). Comparison of progesterone receptor gene expression in TNBC and luminal BC cell lines, normalized expression (TPM) from DepMap 23Q4 Public Dataset (K). Heatmaps report the average of n=3-4 replicates, and use color scales unique to each cell line. Bar graphs report mean ± SD of n=3-4 replicates. Data are reported as cell number relative to vehicle control (VC). Note: cell numbers >100% of VC are shaded as 100% in heatmaps. Conditions that do not share a letter designation are significantly different. See Fig. S4 for comparative viability tables. Welch’s t-test, * p < 0.05, ** p < 0.005, *** p < 0.0005, **** p< 0.0001.

To examine the relative contributions of these non-canonical PRs to TNBC chemoresistance, we implemented the BUP challenge with non-canonical PR-specific agonists and antagonists: the targeted mPR agonist Org OD 02-0, noting that mPR-specific antagonists have not been reported, although MIF interacts with mPRs ^37,43^ (Figure 5A-D); the PGRMC1 antagonist Ag-205 (Figure 5E-H); and the PXR antagonist SPA70 (Figure 5P-S). See viability data tables in Figure S4.

We observed that Org OD 02-0 (ORG) at 10 μM exhibited agonist effects on viability in MDA-MB-231 cells, both on its own (115%) and when combined with 1.0 μM BUP (109%; Figures 5A, B, S4A). However, when administered with 2.0 μM BUP (39.3% viability) or MIF+BUP (43.5% viability), its agonist effects appeared negligible when compared to 2.0 μM BUP alone (40.2% viability) (all p < 0.05;). In T47D cells (Figures 5C, D, S4A), which express *PAQR8* transcripts at a much lower level than MDA-MB-231 cells (Figure 5K), ORG’s agonist effects were less pronounced, although when combined with 2.0 μM BUP or MIF+BUP, cell viability at 30.1% and 29.4% respectively was significantly reduced versus BUP alone at 35.0% (all p < 0.01; Figures 5C, D, S4A). These data suggest mPRβ is likely not a primary driver of chemoresistance in TNBC and luminal BC.

To further investigate the effects of combined PR agonism and antagonism in the background of ORG treatment, we tested BT-20 cells which express mPRβ at a lower level than MDA-MB-231 cells, to avoid the strong agonist effects seen in those cells (Figure S2). We found that ORG alone or added to 0.5 or 1.0 μM BUP did not significantly alter cell survival compared to VC, but including MIF or P4+MIF incurred significant reductions in viable cells versus both concentrations of BUP alone (Figure S2B). Specifically, 0.5 μM BUP alone resulted in 87.7% viability, which was reduced to 59.6% by the addition of ORG+MIF, and further reduced to 49.8% when P4 was included (both p < 0.01). Similarly, 1.0 μM BUP yielded 67.6% viability, and the addition of ORG+MIF elicited 48.3% survival; including P4 in the mix yielded 44.2% (both p < 0.01). While Org+BUP (30.1%) and ORG+MIF+BUP (29.4%) (both p < 0.05) each caused small but significant reductions in cell viability versus BUP alone (35.0%), taken together, these results indicate ORG is not contributing significantly to chemoresistance in TNBC cells in these treatment conditions.

The PGRMC1 antagonist Ag-205 induced a broad pattern of increasing cytotoxicity at higher concentrations in BT-20, HCC1937 and MDA-MB-231 cells, with and without BUP co-treatment (Figure 5E, F; I, J; and L, M respectively; and Figure S4B, C). In BT-20 cells (Figure 5E, F), BUP at 1.0 μM and Ag-205 at 40 μM individually induced significant cell death compared to VC (32.5% and 16.5% viability, respectively; both p < 0.01), and when added together, significantly reduced viability further, to 7.6% (p < 0.05 versus Ag-205 alone). Including 10 μM P4 in this mix reduced surviving cells to 7.0% (p < 0.05 versus Ag-205). To further examine the effects of combined agonism and antagonism when used with 40 μM Ag-205 and 1.0 μM BUP, we added 10 μM MIF to this mix (Figure 5G, H), which significantly reduced cell viability to 1.8% (p < 0.01 when compared to BUP alone). When 10 μM P4 was added, viability was significantly reduced to 2.4% (p < 0.01 versus BUP). While Ag-205+BUP at 1.0 μM reduced the fraction of live cells drastically, to 6.1% versus BUP alone at 28.8% (p < 0.01), adding MIF or P4+MIF was significantly more effective.

In HCC1937 cells (Figure 5I, J, S4B), 2.0 μM BUP and Ag-205 at 40 μM each significantly reduced viable cells versus VC to 55.4% and 53.3% respectively (both p < 0.01; Figure 5I, J), and combining these treatments led to 21.9% viability (p < 0.05 versus Ag-205 alone and p < 0.01 versus BUP alone). Upon the addition of P4, viable cells dropped to 15.3%, a significant reduction from BUP (55.4%; p < 0.01) or Ag-205 alone (53.3%; p < 0.05). We saw a similar pattern in the MDA-MB-231 cells, as Ag-205 alone is more effective than BUP alone, and these cytotoxic effects are enhanced by combining the drugs. Adding in P4, further potentiates the cytotoxic effects. Both 2.0 μM BUP and Ag-204 significantly reduced the fraction of viable cells versus VC, to 71.7% (p < 0.05) and 41.2% (p < 0.01), respectively, with their combination leading to 16.6% viable cells (p < 0.01 compared to BUP alone). The addition of P4 resulted in a significant drop to 6.4% versus BUP and Ag-205 (both p < 0.01). In further examining combined PR agonism and antagonism (Figure 5N, O), we found the addition of MIF to Ag-205+BUP to increase cell survival slightly, from 18.9% for Ag-205+BUP alone to 28.3%, and adding P4 to these treatments likewise led to a slight increase above Ag-205+BUP alone, to 21.4%. However, all of these combinations are significantly more effective than BUP or Ag-205 alone (p < 0.05 for all). In both HCC1937 cells (Figure I, J) and MDA-MB-231 cells (Figure L, M), combining P4 agonism with Ag-205 antagonism and PI3K inhibition provided the strongest reduction in viable cells versus antagonism alone. Together, these findings demonstrate that the non-canonical PGRMC1 is sensitive to simultaneous agonism and antagonism, and can be targeted to enhance PI3K chemotherapeutic effects.

HCC1937 cells express PXR at the highest level in our panel (Figure 5K). At 10 μM, the SPA70 antagonist exhibited a mild effect on cell viability individually (92.9%, ns), which was significantly improved upon combination with BUP at 2.0 μM (51.0%, p < 0.01) (Figures 5P, Q; S4D, E). This effect was significantly enhanced, versus BUP or SPA70 alone, by the addition of 10 μM P4, resulting in 43.4% viability (p < 0.01 for both). We also examined the effects of PR agonism and antagonism on treatment efficacy and found that the addition of MIF to SPA70+BUP led to 30.2% viability versus 42.9% (p < 0.05), a significant reduction (Figures 5 R, S; S4E). Adding P4 to this mix also significantly reduced viability, to 29.9% (p < 0.01) versus SPA70+BUP alone. This data shows that PXR, like PGR and PGRMC1, is a chemotherapeutically targetable PR in TNBC which is vulnerable to simultaneous agonism and antagonism, and can be exploited on its own or in combination with PI3K inhibition to enhance cytotoxic treatment effects.

### Combined progesterone and mifepristone treatment ablates chemoresistant TNBC ALDH1^+^ stem-like cells in response to PI3K inhibition

Given previous findings by ourselves and others on the relationship between BCSC prevalence and chemoresistance in TNBC cell lines ^2,36,75^, we asked whether P4 and/or MIF could influence the prevalence of BCSCs in BUP-treated BT-20 cells. We co-treated the double *PIK3CA*-mutant BT-20 cells with combinations of BUP, P4, and MIF and assayed CSC prevalence 72h later by flow cytometry using both CD44^+^/CD24^-^ status and ALDH1^+^ via the ALDEFLUOR assay, both commonly used markers of BC stem-like cells ^76,77^ (Figure 6A, B). As we observed previously, BUP treatment did not significantly change the prevalence of CD44^+^/CD24^-^ cells, and further P4 and MIF co-treatments exhibited modest effects (Figure 6C, left panel). However, the fraction of CD44^+^/CD24^+^ “drug tolerant persister” cells ^78^ was significantly reduced from 91.1% in VC to 87.7% (p < 0.01) by combined treatment with all three drugs (Figure 6C, right panel). More notably, the ALDH1^+^ CSC cell fraction significantly increased with BUP treatment as we previously reported ^36^, from 0.4% in VC to 2.3% with BUP (p < 0.01; Figure 6D), but this increase was abrogated when P4 and MIF were both combined with BUP treatment, resulting in a decrease to 0.3% (p < 0.01 compared to BUP alone). We subjected the HCC1937 cells to the same treatments (Figure S3), and found that both P4+BUP and MIF+BUP significantly reduced the proportion of ALDH1^+^ cells, from almost 1.0% in VC to 0.3% (p < 0.01) and 0.5% (p < 0.05) respectively (Figure S3D). These results suggest that PR agonist/antagonist perturbations synergistically reduce the prevalence of ALDH1^+^ stem-like cells and CD44^+^/CD24^+^ drug tolerant persister cells in PI3K-targeted TNBC cells.

**Figure 6.**
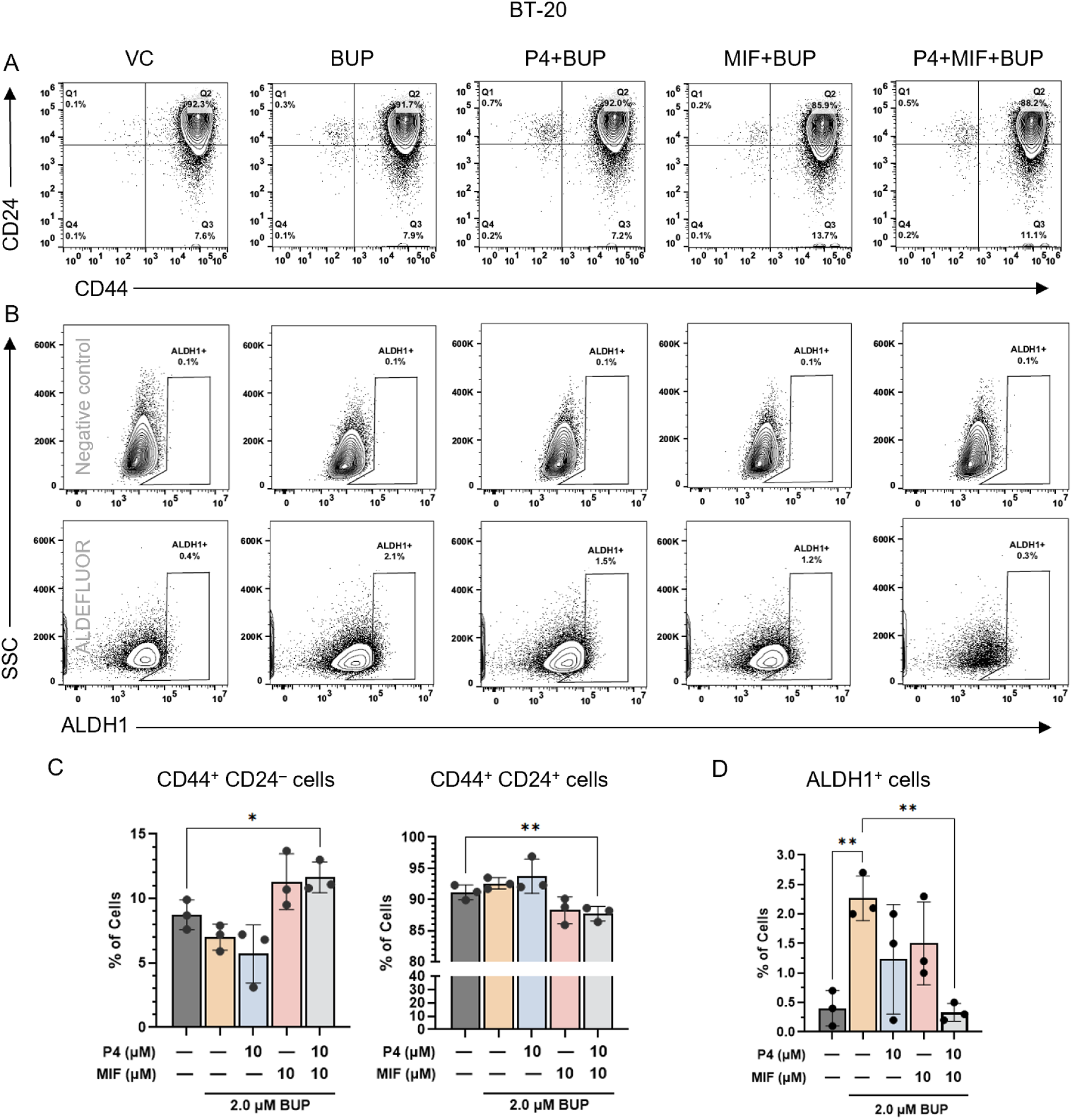
Combined progesterone and mifepristone treatment ablates chemoresistant TNBC ALDH1^+^ stem-like cells in response to PI3K inhibition. Flow cytometric analyses and quantifications of cancer stem-like cell markers in BT-20 TNBC cells treated with vehicle control (VC) or various combinations of progesterone (P4; 10 μM), Mifepristone (MIF; 10 μM), and/or the PI3K inhibitor Buparlisib (BUP; 1 μM) for 72h. Cells were dissociated and incubated with viability dye, anti-CD44 and anti-CD24 antibodies, and the ALDEFLUOR assay, which marks ALDH1^+^ cells. Flow cytometry scatter plots representative of at least 3 replicates in (A-B). Data is mean ± SD in (C-D). ALDELFUOR negative control populations are shown in (B, top row) and were used to draw ALDH1^+^ gates. Quantifications of CD44^+^CD24^+/–^ (C) and ALDH1^+^ cell frequency (D). Welch’s t-test, * p < 0.05, ** p < 0.01.

## Discussion

The effects of P4 on TNBC cells have been largely disregarded since Perou et al. coined the terminology more than 25 years ago ^79^. This framework, however, seems counter-intuitive in the realm of women’s reproductive cancers, and it does not account for the more recent discoveries of canonical and non-canonical PR expression in TNBC ^4–10,80^, and to this list we add p-PGR S345 expression. We found all of these PRs to be functional therapeutic targets in TNBC, capable of inducing cytotoxicity when perturbed with combined agonism and antagonism, with and without concurrent PI3K inhibition (Figures 3, 5). Specifically, simultaneous treatment with P4 and MIF induced significant cell death in all cell lines versus VC, and when combined with PI3K inhibition, also caused significantly more TNBC cell death than BUP alone (Figure 3 C-D, G-H, K-L, O-P). This combinatorial advantage was maintained with an mPR agonist and antagonists targeted to non-canonical PRs as well (Figure 5). Additionally, we found simultaneous PR perturbations and PI3K inhibition to reduce the fraction of ALDH1^+^ BCSCs in TNBC cell lines (Figures 6, S3), a long-standing challenge in the field ^49,50^.

Distinctive functional properties inherent in canonical PGR may underlie the cytotoxicity elicited by simultaneous PR agonism and antagonism. Upon ligand binding, expression of canonical PGR is downregulated ^81–83^; and when bound to MIF in the presence of P4, PGR may form homo- or hetero-dimers and translocate to the cell nucleus, but lose its ability to activate progesterone response elements (PREs) ^84^. In this treatment combination, therefore, PGR functions may be more suppressed than when P4 or MIF alone are present, and this in turn may enhance the effects of PI3K inhibition.

Notably, in addition to the expected agonist effects of P4 and antagonist effects of MIF on cell viability, we also found paradoxical cytotoxicity induced by P4, and enhanced viability elicited by MIF treatment in multiple cell lines (Figure 3A, E and 3B, F, J, respectively). These results may stem from a number of factors including the biphasic effects P4 exerts on CDK2 function and BC cell proliferation ^85–87^, as well as the mixed agonist and antagonist effects of MIF ^88–90^. In addition to concentration-dependent effects, these responses are likely predicated on length of exposure to treatment, the presence of other hormones, tissue type of action ^91^, cell line expression levels of PRs (Figure 5K), and the binding characteristics of MIF ^40,92,93^. While these phenomena may appear intractable, with MIF failing clinical trials against various cancers, for example ^40,94–96^, we ablated MIF’s agonist effects by combining it with the PR agonists P4 (Figure 3) and Org-OD-02 0 (Figures 5A-D, S4A). We see this novel finding as an opportunity to investigate and leverage these mechanisms further. Indeed, using combined PR-targeted agonism and antagonism with PI3K inhibition may enable a reduction in PI3K inhibitor concentrations, reducing toxic and off-target effects for cancer patients.

The cytotoxic advantage in combining PR and PI3K perturbations may arise because these proteins affect a broad range of overlapping signaling and metabolic pathways. For example, PGR and PI3K each promote proliferation in both normal and cancer contexts and are both implicated in nucleotide synthesis ^34,35,97^. Canonical and non-canonical PRs are also thought to enhance PI3K signaling in BC ^7,23,26,28,98^. Similarly, both PGR and PI3K interact with proteins involved in MDR ^99–101^, cell cycle control ^102–104^, and with BRCA1 itself ^105–107^, which is mutated in approximately 15% of TNBC cases ^108^.

The mutational profiles of each cell line may also impact the variations in stress and viability responses to treatment we noted. For example, the two activating *PIK3CA* mutations in the BT-20 cell line sensitize it to PI3K inhibition and may contribute to its most dramatic response of the four cell lines to combined treatments (Figure 3A-D). With only one *PIK3CA* mutation, the T47D cell line exhibited a less stark response (Figure 3M-P). The HCC1937 cell line carries *PTEN* and *BRCA1* mutations which may contribute to its overall lower sensitivity to the three-drug combined treatment than BT-20 and T47D cells (Figure 3E-H). Mutated BRCA1 may allow for greater proliferation in these cells upon exposure to P4, although this effect could be mediated by P4-responsive genes that are differentially expressed between *BRCA1*-mutated and wild type cells ^109,110^. Interestingly, MDA-MB-231 cells, which carry a homozygous mutation in the tumor suppressor gene *CDKN2A*, showed the least sensitivity to either P4 or MIF combined with 2 μM BUP which may indicate this concentration is not sufficient to overcome the likely loss of cell cycle control (Figure 3I-L). MIF is also thought to interact with non-canonical PRs in humans and mice ^37,42,111^, which could in part explain the differing responses to treatment, given that each cell line expresses different levels of mPRβ, PGRMC1 and PXR as described in the DepMap 23Q4 Public Dataset (Figure 5K).

At certain P4 and MIF concentrations, our results comport with the reported mixed effects of P4 on BC ^112–114^ and the cytotoxic potential of MIF ^115^, and MIF combined with chemotherapy ^116^ although in the latter study the authors used lower concentrations of MIF than we, and noted less cell death. The combination of P4 and MIF in TNBC treatment is not well-represented in the literature, although its effects on EMT has been studied ^80^ as has its promotion of apoptosis and effects on cell cycle dynamics in endometrial and cervical cancer cells ^117^.

Previous findings that BC treatment with chemotherapy can increase the BCSC fraction ^49,118,119^ was borne out in our studies, with BUP treatment resulting in a significantly increased proportion of ALDH1^+^ BCSCs in BT-20 cells (Figure 6). Strikingly, however, we found that combined BUP, P4 and MIF treatment decrease the ALDH1^+^ BCSC fraction more than BUP alone in this context, and BUP+P4 reduced this fraction below that of VC in HCC1937 cells (Figure S3), which could point to a promising advantage in these combinatorial treatments. Since ALDH1 expressing cells are considered enriched in TNBC ^2^ and ALDH1 is negatively correlated with PGR expression in neoplastic breast tissue ^120^, the basis of our observed effects suggests further investigation into the interactions between these two proteins is warranted.

While our results clearly indicate therapeutic potential in combined PR and PI3K perturbation in TNBC, we recognize some limitations in our investigation. Some of our treatments may effect off-target responses, including MIF’s modulation of glucocorticoid receptors and Ag-205’s actions on actin-associated proteins and lipid synthesis ^46,121^, which may necessitate further treatment optimization. Longitudinal studies including scRNA-seq could provide insight into the genomic effects of canonical and non-canonical PR modulation in TNBC alone and in combination with PI3K inhibition, possibly pointing to additional targets or the source of off-target effects. Genomic studies could aid in the identification of affected downstream P4 targets which may lend themselves to PR-sparing treatments. Additionally, PR overexpression or chemical or siRNA knockdown could shed light on each receptor’s role in resistance and the utility of targeting one or more receptors. These treatments did not ablate CD44^+^/CD24^-^ or ALDH1^+^ BCSCs, though greater efficacy may have been obtained with different treatment durations, concentrations, or repeated or temporally-sequenced administration. While *BRCA1*, *CDKN2A* and *PIK3* mutations are well-studied, whether these TNBC cell lines harbor mutations in any PRs, and the effects of these mutations on PR function and chemoresistance should also be investigated. Continuation of this work in syngeneic or genetically modified BC mouse models, and the use of PDX tissue would provide a fuller picture of the physiological relevance of our work and provide insight into safe and effective dosages. Given P4’s pleiotropic roles in mammalian homeostasis ^122,123^, isolating PR modulating treatments to tumor tissue is of utmost importance, using state of the art nanoparticles, or antibody or RNA conjugation.

Lastly, even in menopause, the effects of P4 on reproductive tissues ostensibly should not be disregarded, not only for the possibility of non-canonical PR expression, but sensitive cells are equipped with the biochemical machinery to synthesize steroids in an intracrine fashion ^124–126^, thus malignancies or hormonal imbalances may be fueled intrinsically within each cell. Notably, this phenomenon may underlie the aggressiveness of castration-resistant prostate cancer^127,128^.

We have much to learn about P4 and PR functions in TNBC; indeed P4-responsive proteins are still being identified in female reproductive tissue, as was ABHD2 in 2016 ^129,130^. We propose a new approach in designing TNBC-targeted chemotherapies which include PGR, mPRs, PGRMC1 and/or PXR perturbations, and a reconsideration of the classification of this aggressive disease.

## Materials and Methods

### Cell Culture

Cells were maintained in two-dimensional culture at 5 x 10^4^ cells/mL and 37°C in 5% CO_2_ in a humidified chamber. BT-20 cells were cultured in growth medium using Minimum Essential Medium (MEM; Corning, Inc., Oneonta, NY, #10-009-CV) with 10% fetal bovine serum (FBS; Corning, Inc., #35-010-CV) and 1% penicillin/streptomycin (ThermoFisher, #15140122). Growth medium for both HCC1937 and MDA-MB-231 cells was Roswell Park Memorial Institute 1640 (RPMI 1640; Corning, Inc., #10-041-CVR) with 10% FBS and 1% penicillin/streptomycin. T47D cells were grown in RPMI supplemented with 10% fetal bovine serum, 1% penicillin/streptomycin and 0.2 Units/mL bovine insulin (Millipore Sigma, #I-035). BT-20 and HCC1937 cells were also cultured in charcoal-stripped FBS (Gibco, #12676-029) and phenol red-free MEM (Corning, #17-305) and RPMI respectively (Corning, #17-105). Phenol red-free MEM was supplemented with L-glutamine (Corning, #25-005-CI), sodium pyruvate (Corning, #25-000-CIR) and NEAAs (Gibco, #11140-050); and phenol red-free RPMI was supplemented with L-glutamine (Corning, #25-005-CI) and HEPES (Gibco, #15630-080). Cancer cell lines were passaged fewer than 20 times. MDA-MB-231 and T47D cells were generous gifts from Claudia Fischbach-Teschl and Jan Lammerding, respectively, of Cornell University. Other cell lines were purchased from American Type Culture Collection (atcc.org, Manassas, VA).

### Cell Viability

Progesterone, mifepristone and Buparlisib were obtained from Selleck Chemicals LLC (Houston, TX, #S1705, #S2606 and #S2247 respectively) and reconstituted in DMSO (Sigma-Aldrich, #D8418). Ag-205 and SPA-70 was obtained from Sigma-Aldrich (#A-1487 and #SMA-2662 respectively), and each was dissolved in DMSO at stock concentrations. Cells were plated at a density of 5 x 10^4^ cells/mL and allowed to attach for 24h, then treated with progesterone, mifepristone, and/or Buparlisib at the indicated concentrations up to 72h. Vehicle control cells were maintained in normal growth medium supplemented with 0.1% DMSO, and DMSO was added to inhibitor-treated cells to normalize its concentration. Cells were not included in in 96-well plate edge wells to minimize edge effects. Methanol fixation was followed by propidium iodide staining (ThermoFisher, #P3566), cells were counted using CellProfiler (Broad Institute, Cambridge, MA) ^131^, and data was normalized to vehicle control cells. MATLAB (MathWorks, Natick, MA) and Prism (GraphPad Software, Boston, MA) were used for heatmap graphs, and Prism and Excel were used for statistical analyses.

### Imaging

Nikon Eclipse Ti-E Microscope (MVI, Avon, MA) with Spectra X or Celesta light sources was used to capture digital images. For viability studies, propidium iodide was used to stain nuclei; images were captured in the TRITC channel; thresholding performed in native NIS Elements software; nuclear segmentation and automated counting were performed using CellProfiler.

### Immunocytochemistry

Cells were fixed in 4% formaldehyde for 12 mins at room temperature, followed by three 5 min PBS washes. Cells were then permeabilized with either 0.3% TritonX-100 or 0.5% Tween in PBS for 20min, depending on cellular compartment of target protein. Donkey serum at 10% in either 0.01% TritonX-100 or 0.1% Tween in PBS was used to block for 20 mins at RT. Primary and secondary antibodies were diluted in PBS containing 1% donkey serum and either 0.01% TritonX-100 or 0.1% Tween in PBS. Primary antibodies were incubated overnight at 4°C with gentle agitation. Secondary staining was performed for 1h at room temperature. Primary antibodies used from Invitrogen: α-Tubulin (Rat mAb, clone YL1/2, #MA1-80017); Phospho-PR (Ser345, rabbit pAb, #PA5-106027), PXR (rabbit pAb, #PA5-72551); PR (rabbit mAb, clone R.380.2; #MA5-14841). Cell Signaling Technology primary antibodies: PGRMC1 (clone D6M5M, rabbit mAb, #13856); PR-A/B (clone D8Q2J, rabbit mAb, #8757). Also used were: mPRβ (Bioss USA, Woburn, MA, rabbit pAb, #bs-11410R; and Abcepta, San Diego, CA, rabbit pAb, #AP14849b). Secondary antibodies were from Invitrogen: Goat anti-rabbit/AF 647, #A21245 and goat ant-rat/AF555, #A21434. DAPI was used for nuclear staining (Invitrogen #D3571). Positively-stained cells were counted with CellProfiler where possible, or manually. Nuclear staining of PGR and p-PGR S345 were counted per nucleus; all others by cell body.

### Quantitative Immunoblotting

After appropriate treatments, cells were lysed with RIPA or NP-40 buffer as appropriate for cellular compartments of target proteins. RIPA: 50 mM Tris-HCl, 150 mM NaCl, 1% Triton X-100, 0.5% sodium deoxycholate, 0.1% SDS, 5 mM EDTA supplemented with 10 μg/mL aprotinin, 10 μg/mL leupeptin, 1 μg/mL pepstatin, 1 mM PMSF, 1 μg/mL microcystin-LR, 200 μM sodium orthovanadate. NP-40: 50 mM Tris-HCl, 150 mM NaCl, 0.5% NP-40 substitute, 5 mM EDTA supplemented as above. Cells were placed on ice and washed with cold PBS. Cold lysis buffer was added to each well and allowed to incubate on ice for 15 minutes. Cells were scraped and lysates were collected, then centrifuged at 4° C for 10 minutes at 15,000 x g. The supernatant was collected and protein concentration was quantified with a micro bicinchoninic acid assay kit (ThemoFisher, Waltham, MA, #23235). Lysates were electrophoresed and transferred using the ThermoFisher BOLT Bis-Tris system (#NW0008C). Each lysate was loaded with Bolt sample buffer and reducing agent onto electrophoresis gels at 10 - 20 μg of protein per lane, with chemiluminescent protein standards (Bio-Rad, Hercules, CA, #1610374 and #1610376) per manufacturer’s recommendations. Gels were run using Bolt MES running buffer at 200 V for 35 minutes. Proteins were wet-transferred at RT to a methanol-activated PVDF membrane using BOLT transfer buffer, methanol and anti-oxidant at 20 V for 60 minutes. Membranes were blocked in 5% bovine serum albumin (BSA) (Rockland Immunochemicals, #BSA-50) in Tris-buffered saline with Tween (TBST) for 1h with gentle rocking at RT. Primary antibodies were diluted in 5% BSA/TBST at the following concentrations: mPRβ (Bioss USA, rabbit pAb, #bs-11410R; and Abcepta, 40 kDa, rabbit pAb, #AP14849b), 1:1000. All following are ThermoFisher/Invitrogen: GAPDH (36 kDa, rabbit polyclonal Ab, #PA1-987), 1:10,000; Phospho-PR (Ser345, 94, 120 kDa, rabbit pAb, #PA5-106027), 1:1000; PXR (rabbit pAb, #PA5-72551), 1:1000; PR (94, 120 kDa, rabbit mAb, clone R.380.2; #MA5-14841), 1:1000. All following are Cell Signaling Technology (Danvers, MA): α-Actinin (100 kDa, rabbit mAb, clone D6F6, #6487), 1:1000; Cleaved PARP (Asp214, 89 kDa, rabbit mAb, #9541), 1:1000; Phospho-p38 (Thr180/Tyr192, 43 kDa, clone D3F9, rabbit mAb, #4511), 1:1000; Phospho-p44/42 MAPK (ERK 1/2) (Thr202/Tyr204, 42, 44 kDa, clone D13.14.4E, rabbit mAb, #4370), 1:2000; PGRMC1 (25 kDa, clone D6M5M, rabbit mAb, #13856), 1:1000; Phospho-Akt (Ser473, 60 kDa, clone D9E, rabbit mAb, #4060), 1:2000; Phospho-PR (Ser345, 90, 118 kDa; #12783), 1:1000; PR-A/B; 90, 118 kDa, clone D8Q2J, rabbit mAb, #8757), 1:1000; PXR (45 kDa, clone E4S1W, rabbit mAb, #44646), 1:1000; Vinculin (124 kDa, rabbit mAb, clone E1E9V, #13901), 1:1000. Blots were incubated with primary antibodies with gentle rocking at 4° C overnight, then washed with TBST three times for five minutes each. Blots were then incubated with goat anti-rabbit secondary antibody HRP conjugate (Jackson ImmunoResearch Laboratories, West Grove, PA, #111-005-045, 1:5,000 dilution) and HRP-Streptavidin Conjugate (Bio-Rad, #1610380) at 1:10,000 dilution in TBST for 1h at RT, then washed again as above. Blots were then incubated in ECL substrate (Bio-Rad, #1705060) for 1 minute at RT and imaged on the ChemiDoc chemiluminescent system (Bio-Rad, #17001401). Protein expression was normalized to housekeeping genes and quantified using ImageJ, Excel and GraphPad Prism software.

### Flow cytometry

Briefly, for CD44 and CD24 assays, cells were harvested with Accutase (Innovative Cell Technologies, Inc., San Diego, CA, #AT104) to preserve surface proteins, then treated with FC block (ThermoFisher, #14-9191-73) before staining with BioLegend (San Diego, CA) antibodies as follows: anti-CD44 antibody conjugated to BV605 (clone IM7, #103047), and anti-CD24/AF647 (clone ML5, #311110). DAPI (ThermoFisher, #D3571), was used for live/dead cell discrimination.

For ALDH1 assays, the ALDEFLUOR kit was used from STEMCELL Technologies, Inc. (Vancouver, BC, Canada, #1702), per the manufacturer’s protocol.

Flow cytometry was performed on the Sony MA900 flow cytometer (Sony Biotechnology, San Jose, CA) in Cornell University’s Biotechnology core. Flow data analysis performed using FlowJo software (Becton Dickinson and Company).

## Statistical analyses

Unless stated otherwise, experimental data are calculated as mean values normalized to vehicle control data, with error bars representing ±SD for a minimum of three biological replicates. Differences in means between experimental groups was evaluated using Welch’s t-tests, and a p-value of less than 0.05 was considered significant. GraphPad Prism, MATLAB and Microsoft Excel were used for data analysis and visualization.

## Conflicts of interest

The authors declare no conflicts of interest.

### Author contributions

P.E.P. conceived of the work, conducted all experiments and analyses, and wrote the manuscript. J.W.C. also conducted some experiments. B.D.C. conceived of the work and wrote the manuscript.

## Acknowledgments

The authors would like to thank Joseph Druso of the Fischbach-Teschl lab for assistance in culturing MDA-MB-231 cells; and Marc Antonyak of the Cerione lab for assistance in Western blotting. We extend our thanks to Lydia Tesfa, Jaclyn Mahoney and Michael Sledziona at the Flow Cytometry Facility (RRID:SCR_021740) of the Cornell Biotechnology Resource Center for their help in performing flow cytometry experiments. This work was supported by the US National Institutes of Health (NIH) grants U54CA210184, R01CA238745, R01CA260115 and R01CA248524 (B.D.C.) and Cornell Stem Cell Training Fellowship (P.E.P.). The content is solely the responsibility of the authors and does not necessarily represent the official views of the NIH.

---

**Figure S1.**
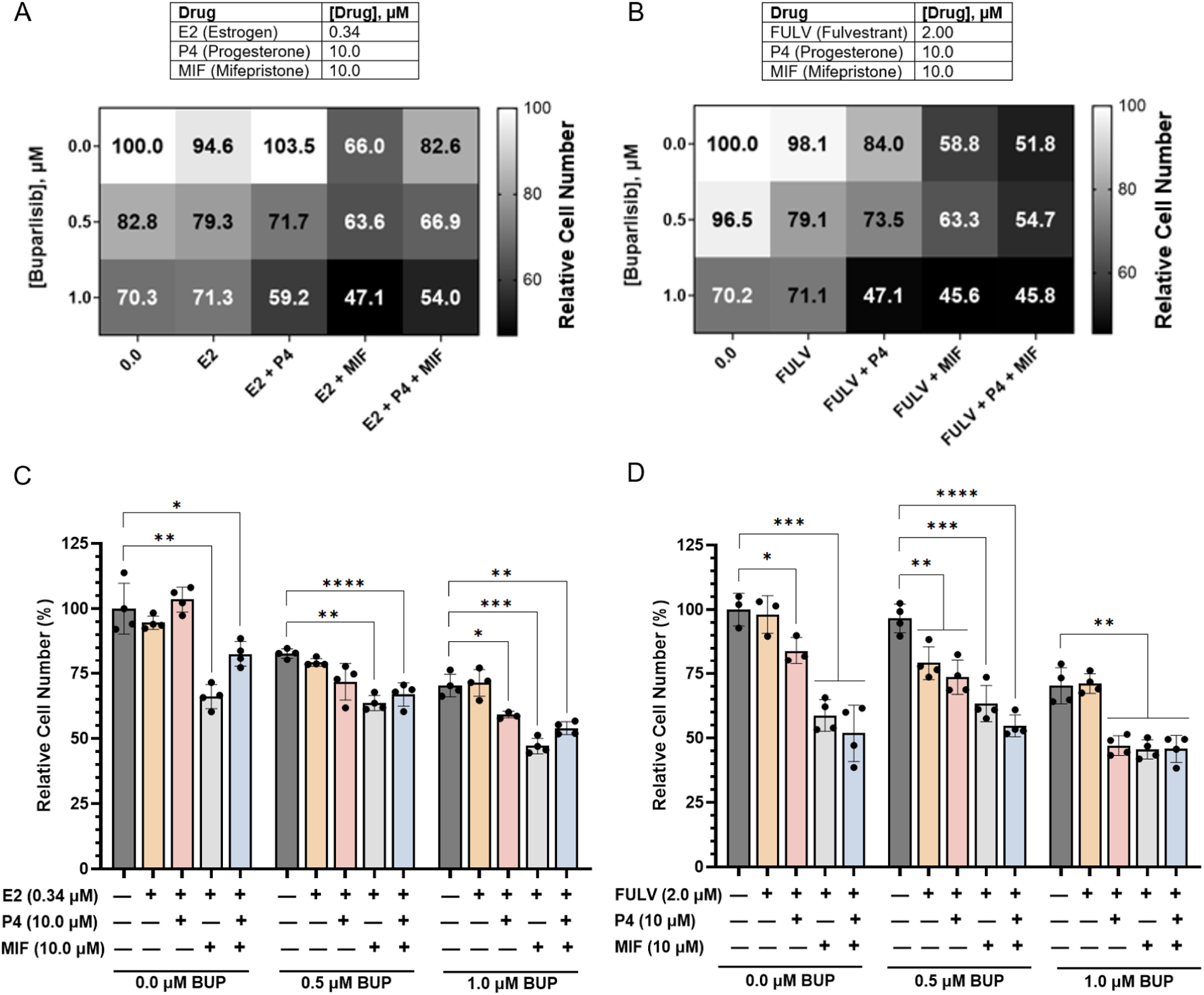
Simultaneous perturbation of estrogen and progesterone receptors restricts TNBC cell viability when combined with PI3K inhibition. BT-20 TNBC cells were treated for 72h with progesterone (P4; 10 μM), estrogen (E2; 0.34 μM); Mifepristone (MIF; 10 μM), the estrogen receptor antagonist fulvestrant (FULV; 2 μM), and/or the PI3K inhibitor Buparlisib (0-1 μM) in various combinations, and relative cell viability was measured. Cell numbers >100% of VC are shaded as 100% in heatmaps. Data is reported as cell number relative to vehicle control (VC), mean of n=3+ replicates. Bar graphs depict mean ± SD; Welch’s t-test; **** p ≤ 0.0001; *** p ≤ 0.0005; ** p < 0.005; * p < 0.05.

**Figure S2.**
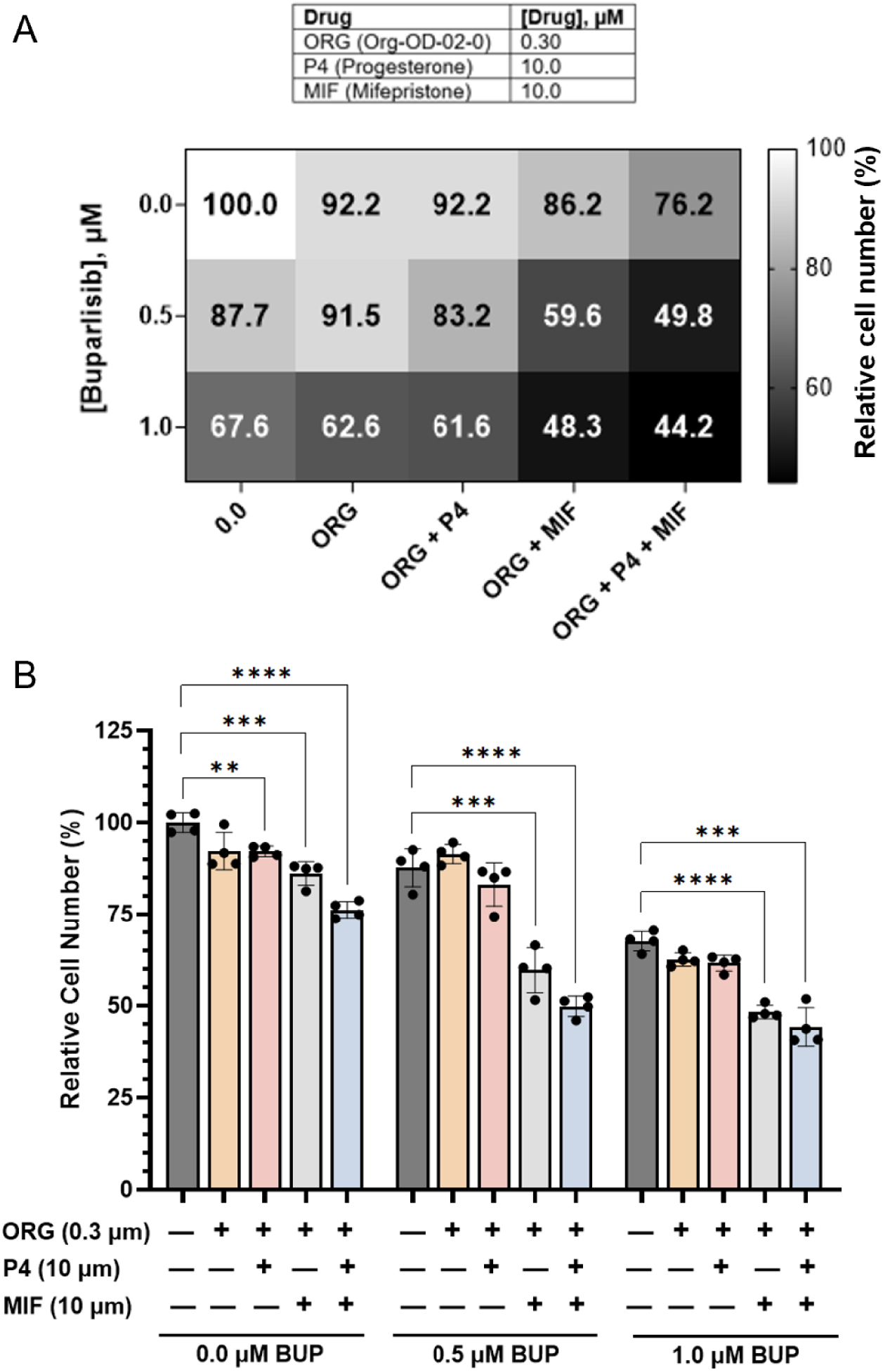
Combinatorial perturbation of mPRβ receptor signaling restricts TNBC cell viability when combined with mifepristone and/or PI3K inhibition. BT-20 TNBC cells were treated for 72h with mPR agonist Org OD 02-0 (ORG) at 0.3 μM, PI3K inhibitor Buparlisib (BUP) at 0-1 μM, progesterone (P4) 10 μM, and/or mifepristone (MIF) 10 μM in various combinations, and relative cell viability was measured (A). Data are reported as cell number relative to vehicle control (VC); mean of n=3+ replicates. Bar graphs (B) depict mean ± SD; Welch’s t-test; **** p ≤ 0.0001; *** p ≤ 0.0005; ** p < 0.005; * p < 0.05.

**Figure S3.**
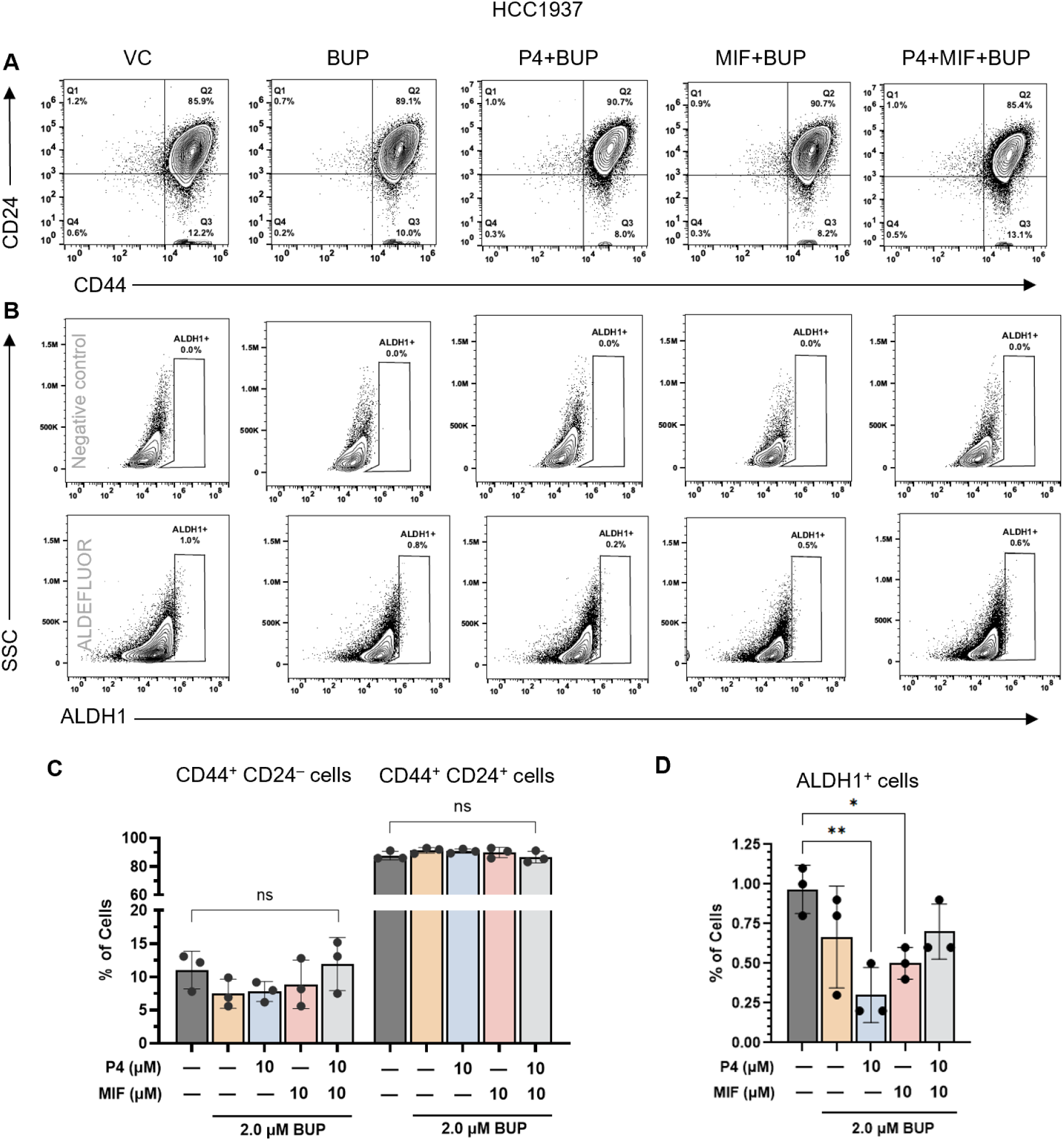
Combined progesterone and Buparlisib treatment ablates chemoresistant TNBC ALDH1^+^ stem-like cells. Flow cytometric analyses and quantifications of cancer stem-like cell markers in HCC1937 TNBC cells treated with vehicle control (VC) or various combinations of progesterone (P4; 10 μM), Mifepristone (MIF; 10 μM), and/or the PI3K inhibitor Buparlisib (BUP; 1 μM) for 72h. Cells were dissociated and incubated with viability dye, anti-CD44 and anti-CD24 antibodies, and the ALDEFLUOR assay, which marks ALDH1^+^ cells. Flow cytometry scatter plots representative of at least 3 replicates in (A-B). Data is mean ± SD in (C-D). ALDELFUOR negative control populations are shown in (B, top row) and were used to draw ALDH1^+^ gates. Quantifications of CD44^+^CD24^+/–^ (C) and ALDH1^+^ cell frequency (D). Welch’s t-test, * p<0.05, ** p<0.01.

**Figure S4.**
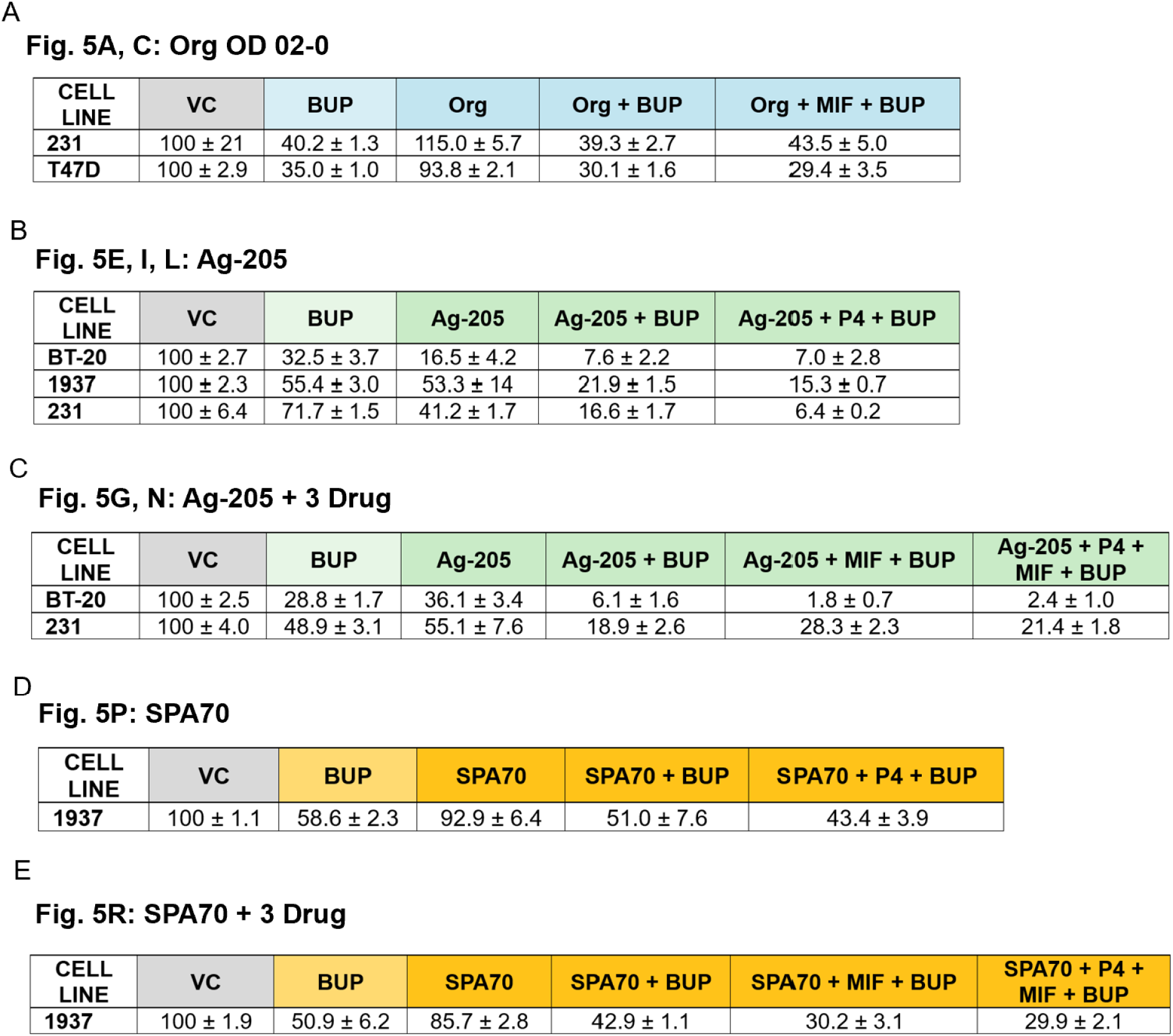
Comparative viability data tables for. Figure 5. TNBC cells BT-20, HCC1937 (1937), MDA-MB-231 (231) and luminal A cell line T47D were treated for 72h with various combinations of mPR agonist Org OD 02-0 (ORG) at 10 μM (A); PGRMC1 antagonist Ag-205 at 40 μM (B, C); PXR antagonist SPA70 at 10 μM (D, E); and/or progesterone (P4) at 10 μM (B-E); mifepristone (MIF) 10 μM (A, C, E); and PI3K inhibitor Buparlisib (BUP) at 1.0 μM in BT-20 cells and 2.0 μM in all others (A-E), and relative cell viability was measured. Data are reported as % cell viability normalized to vehicle control (VC) ± SD; n=3-4 replicates.

